# A stutter in the coiled coil domain of *E.coli* co-chaperone GrpE connects structure with function

**DOI:** 10.1101/2020.07.03.186056

**Authors:** Tulsi Upadhyay, Vaibhav V Karekar, Ishu Saraogi

## Abstract

In bacteria, the co-chaperone GrpE acts as a nucleotide exchange factor and plays an important role in controlling the chaperone cycle of DnaK. The functional form of GrpE is an asymmetric dimer, consisting of a long non-ideal coiled-coil. During heat stress, this region partially unfolds and prevents DnaK nucleotide exchange, ultimately ceasing the chaperone cycle. In this study, we elucidate the role of thermal unfolding of the coiled-coil domain of *E. coli* GrpE in regulating its co-chaperonic activity. The presence of a stutter disrupts the regular heptad arrangement typically found in an ideal coiled coil resulting in structural distortion. Introduction of hydrophobic residues at the stutter altered the structural stability of the coiled-coil. Using an in vitro FRET assay, we show for the first time that the enhanced stability of GrpE resulted in an increased affinity for DnaK. However, the mutants were defective in *in vitro* functional assays, and were unable to support bacterial growth at heat shock temperature in a *grpE*-deleted *E. coli* strain. This work provides valuable insights into the functional role of a stutter in the GrpE coiled-coil, and its role in regulating the DnaK-chaperone cycle for bacterial survival during heat stress. More generally, our findings illustrate how a sequence specific stutter in a coiled-coil domain regulates the structure function trade-off in proteins.

## Introduction

Molecular chaperones like Hsp70 (Heat Shock Protein-70) play an active role in maintaining protein homeostasis in living organisms. The Hsp70 chaperone system is involved in folding of newly synthesized proteins, preventing protein misfolding, assisting in disaggregation, and transportation of proteins through the membrane^1,2^. In bacteria, this chaperone machinery consists of a homologue of Hsp70, known as DnaK. For its efficient functioning, two co-chaperones DnaJ and GrpE synergistically stimulate the ATP-dependent activity of DnaK^3–7^. The co-chaperone GrpE is a nucleotide exchange factor that is essential for bacterial survival, and also acts as a thermosensor^8–11^. It regulates the bacterial Hsp70 cycle by switching the function of DnaK from its normal foldase state to a holdase state during stress^12,13^. Under physiological conditions, GrpE acts as a nucleotide exchange factor to exchange ADP with ATP in the DnaK nucleotide-binding domain^14,15^. This nucleotide exchange maintains the exact balance between low and high-affinity states of DnaK for its protein substrate, and controls the foldase activity of DnaK (Fig.1a)^16,17^. During heat stress, GrpE experiences a significant structural change, which alters its interaction with DnaK, which in turn activates the holdase function of DnaK (Fig.1a)^12,16^.

**Fig. 1.**
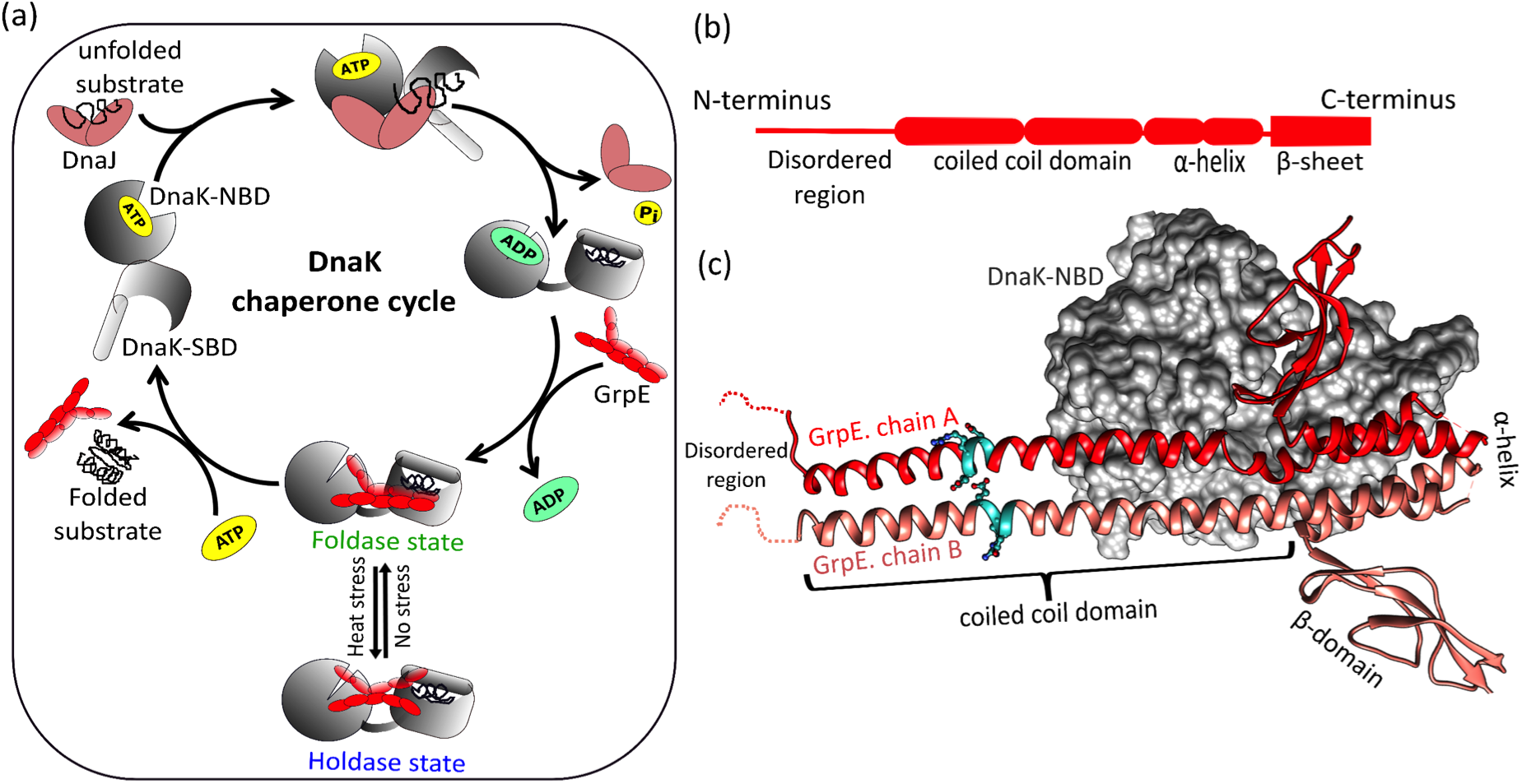
The structure of co-chaperone GrpE. (a) The DnaK chaperone cycle in bacteria. Under normal conditions, DnaK (grey) interacts with co-chaperone DnaJ (brown) and GrpE (red) to maintain protein homeostasis. During heat stress, structural changes in GrpE halt the chaperonic cycle preventing futile ATP hydrolysis (b) GrpE can be roughly divided into four regions: an N-terminal disordered region, a coiled-coil domain, an α-helical region, and a β-sheet domain (c) Co-crystal structure of DnaK nucleotide binding domain (DnaK-NBD) complexed with GrpE dimer (PDB ID:1DKG). Only chain A of GrpE (red) is involved in interaction with DnaK (grey surface). A stutter in the coiled-coil domain is represented in cyan colour. The image was generated with UCSF Chimera^7,18^.

The co-crystal structure of GrpE with the nucleotide-binding domain of DnaK (DnaK-NBD) shows an asymmetric GrpE dimer, where only one chain (labeled chain A) of the co-chaperone interacts with DnaK (Fig.1c)^18^. The GrpE dimer is structurally divided into four parts: the N-terminal unstructured region (residues 1-33), which is involved in substrate release from DnaK; the extended coiled-coil region (80 Å, residues 34-88), that plays an important role in its thermosensing activity; and a 4-helical bundle (residues 89-137) followed by a β-sheet wing region (residues 137-197) which are involved in nucleotide exchange during the chaperone cycle (Fig.1b) ^8,19,20^. During heat stress, the coiled coil domain of GrpE interacts with substrate binding domain of DnaK (DnaK-SBD) and stabilizes the DnaK-substrate complex^16,21^. However, it is not known whether structural rearrangements due to stutter in the coiled-coil domain of GrpE play a role in modulating its interaction with DnaK, and in turn its co-chaperonic activity.

Coiled-coil domains are common structural motifs present in various globular proteins and show diverse functions like assisting in proteins oligomerization, providing mechanical and thermal stability, and facilitating protein-protein interactions^22,23^. Coiled coils are also found in molecular chaperones, where they regulate protein-protein interactions and ensure proper protein homeostasis^24,25^. Some coiled-coils in bacteria have been reported to play a significant role in virulence and host-pathogen interactions^26,27^. Coiled-coils are composed of heptad repeats assigned as [a-b-c-d-e-f-g]_n_. The ‘a’ and ‘d’ positions of two heptads facing each other have hydrophobic amino acids, and the ‘e’ and ‘g’ positions consist of charged residues that are involved in interhelical ion pairing^28–30^. This amino acid arrangement contributes to the stability of coiled-coil by forming ‘knobs-into-holes’ packing. This packing is disrupted by addition of destabilizing residues (charged residues or alanine) at the ‘a’ and ‘d’ positions. Other types of alterations in the heptad repeats may include one to two extra residues referred to as ‘insertions’, or three to four residues referred to as ‘stammer’ and ‘stutter’, respectively^31^. These changes in the coiled coil domain convert the ‘knobs-into-holes’ packing into ‘knobs-into-knobs’ packing, which changes the degree of supercoiling in the coiled-coil domain. This often leads to progressive delocalization of the periodicity of the residues, which modulates the interhelical interactions and changes the stability of the coiled-coil region^22,32^.

The coiled-coil domain of GrpE plays an essential role in detecting changes in ambient temperature during bacterial growth. These changes are thought to be sensed by thermal unfolding of *E. coli* GrpE, which displays two well-separated thermal transitions with midpoints at ~50°C and ~75°C^9^. The first transition has been attributed to the unfolding of the coiled-coil domain, whereas the second transition is likely due to unfolding of the four helical bundle domain^33^. In this work, we elucidate the mechanism by which information about molecular changes in the GrpE coiled-coil domain is transmitted to the DnaK binding domain, ultimately leading to control of GrpE co-chaperonic function.

The GrpE coiled-coil domain contains two stutters (i + 3, i + 4, i + 3, i + 4, **<u>i + 4</u>**, i + 3, i + 4, i + 3, i + 4, **<u>i + 4</u>**) forming two simultaneous heptad (i +3, i + 4) and hendecad (i + 3, i + 4, **<u>i + 4</u>**) repeats (Fig. 2a)^18,34^. The first stutter ‘ERDG’ in the coiled-coil domain starts at residue 58 and ends at residue 61 (Fig.1b & Fig. 2a), while the second stutter ‘TELD’ is located at residues 76 to 79. In this work, using various biophysical and biochemical tools, we report that an increase in hydrophobicity in the first stutter region changes the structure and stability of the coiled-coil domain. Using an *in vitro* FRET assay, we show for the first time that mutations in the coiled-coil domain lead to a significant change in the DnaK-GrpE binding interaction. However, the mutants are found to be defective in *in vitro* functional assays and bacterial growth assays, providing valuable insights into the functional role of a stutter in the GrpE coiled-coil, and its role in regulating the DnaK-chaperone cycle for bacterial survival during heat stress.

**Fig. 2.**
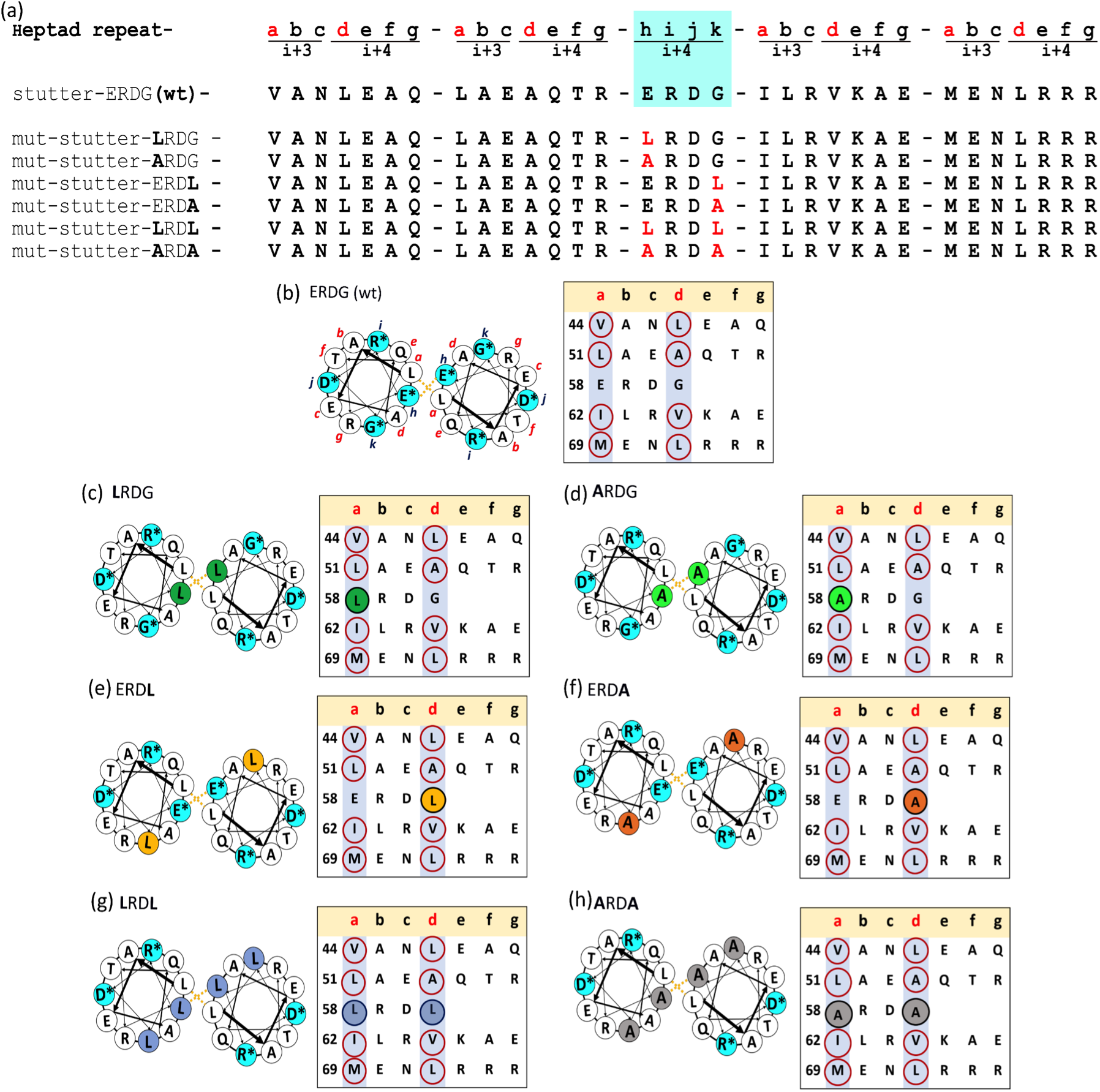
The coiled-coil domain of co-chaperone GrpE. (a) The first row shows typical heptad repeats [a, b, c, d, e, f, g] with stutter [h, i, j, k] insertion. Wild-type *E. coli* GrpE coiled coil domain and the mutants used in this study are listed. (b) Helical wheel diagram for two hendecad repeats showing interhelical interaction (left panel). Stutter residues are in cyan. Yellow dotted lines show hydrophobic interaction between residues at positions a, d, h, and e. Tabular arrangement of the coiled-coil domain residues showing the heptad repeats and the stutter positions for ERDG (wt) (right panel). (c)-(h) Corresponding arrangement for all the single and double stutter mutants used in this study are shown (c) **L**RDG (green) and (d) **A**RDG (light green), (e) ERD**L** (yellow), (f) ERD**A** (brown) (g) **L**RD**L** (blue) and (h) **A**RD**A** (grey). The residue positions were determined using the software COILS^35^.

## Results

### Introduction of hydrophobic residues in the stutter increases the stability of the GrpE mutants

We first mapped the position of the residues in the heptad repeats of the GrpE coiled-coil domain using the software COILS (from ExPaSy-Bioinformatics tool portal)^35^, and identified the stutter forming residues in the sequence (Fig.2a). The helical wheel diagram was different from the canonical heptad repeats, and was arranged in the hendecad repeat [a, b, c, d, e, f, g, **h, i, j, k**]_n_ for the two monomers. Here, a, e, h, and d residues present in the core were involved in hydrophobic interactions, and i and k residues might engage in electrostatic interactions depending on the sequence (Fig. 2b)^36^. We observed that the first residue of the stutter ‘ERDG’, with ‘E’ (-vely charged residue at the 58^th^ position) occupied the ‘h’ position of the stutter and faced the core; and the last residue ‘G’ (neutral residue at the 61^st^ position) at the ‘k’ position faced the hydrophilic region of the coiled-coil domain (Fig. 2b). The assignment of these two destabilizing residues to the ‘h’ and ‘k’ positions was supported by a visual analysis of the GrpE-DnaK co-crystal structure (PDB ID: 1DKG)^18^.

We reasoned that the presence of destabilizing residues in the interhelical hydrophobic core may alter the heptad periodicity of the coiled-coil domain, leading to a distorted coiled coil. We examined the role of ‘E’ and ‘G’ in the stutter of wt-GrpE (ERDG) by substituting these two positions with either a highly hydrophobic amino acid leucine (Leu, L) or with a neutral amino acid alanine (Ala, A) or both. According to the coiled coil periodic table, Leu occurs most frequently at the core of coiled-coil domains and acts as a stabilizing residue^37,38^. On the other hand, Ala has a high α-helical propensity, which provides stability to the core and is expected to increase the α-helical content^39^. We hypothesized that introduction of Leu or Ala might make the coiled coil core more compact, and this packing effect might be propagated towards the neighbouring interhelical region thereby modulating the overall coiled coil stability. To probe this hypothesis, we designed six mutants; two mutants at position 58 henceforth referred to as **L**RDG and **A**RDG, two mutants at position 61 referred to as ERD**L** and ERD**A,** and two double mutants referred to as **L**RD**L** and **A**RD**A** (Fig. 2a and c-h).

Using far-UV circular dichroism (CD) spectroscopy, we measured changes in the secondary structure of the stutter mutants and compared them to wild type GrpE. The mutants with substitution at the 58^th^ position i.e. **L**RDG and **A**RDG showed a large increase in helicity (Fig. 3a). In contrast, single stutter mutants with substitution at 61^st^ position ERD**L** and ERD**A**, and double stutter mutants **L**RD**L** and **A**RD**A** showed no significant change in helicity (Fig. 3b,c). No significant change was seen in the θ_222_/θ_208_ ratio compared to ERDG (wt) confirming that the global structure of the mutant proteins remained unchanged (Table 1)^40^.

**Table 1.**
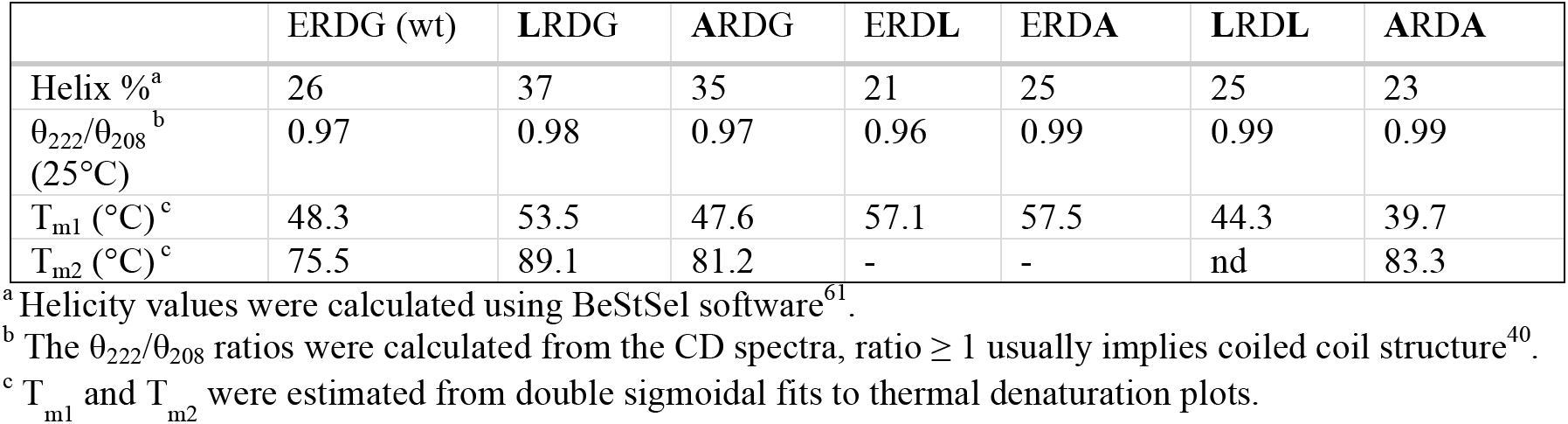
Helicity ratios (θ_222_/θ_208_) and thermal melting transitions for wt-GrpE and stutter mutants.

|  | ERDG (wt) | LRDG | ARDG | ERDL | ERDA | LRDL | ARDA |
| --- | --- | --- | --- | --- | --- | --- | --- |
| Helix %<sup>a</sup> | 26 | 37 | 35 | 21 | 25 | 25 | 23 |
| $\theta_{222}/\theta_{208}$<sup>b</sup><br>(25°C) | 0.97 | 0.98 | 0.97 | 0.96 | 0.99 | 0.99 | 0.99 |
| $T_{m1}$ (°C)<sup>c</sup> | 48.3 | 53.5 | 47.6 | 57.1 | 57.5 | 44.3 | 39.7 |
| $T_{m2}$ (°C)<sup>c</sup> | 75.5 | 89.1 | 81.2 | - | - | nd | 83.3 |

<sup>a</sup> Helicity values were calculated using BeStSel software<sup>61</sup>.
<sup>b</sup> The $\theta_{222}/\theta_{208}$ ratios were calculated from the CD spectra, ratio $\geq 1$ usually implies coiled coil structure<sup>40</sup>.
<sup>c</sup> $T_{m1}$ and $T_{m2}$ were estimated from double sigmoidal fits to thermal denaturation plots.

**Fig. 3.**
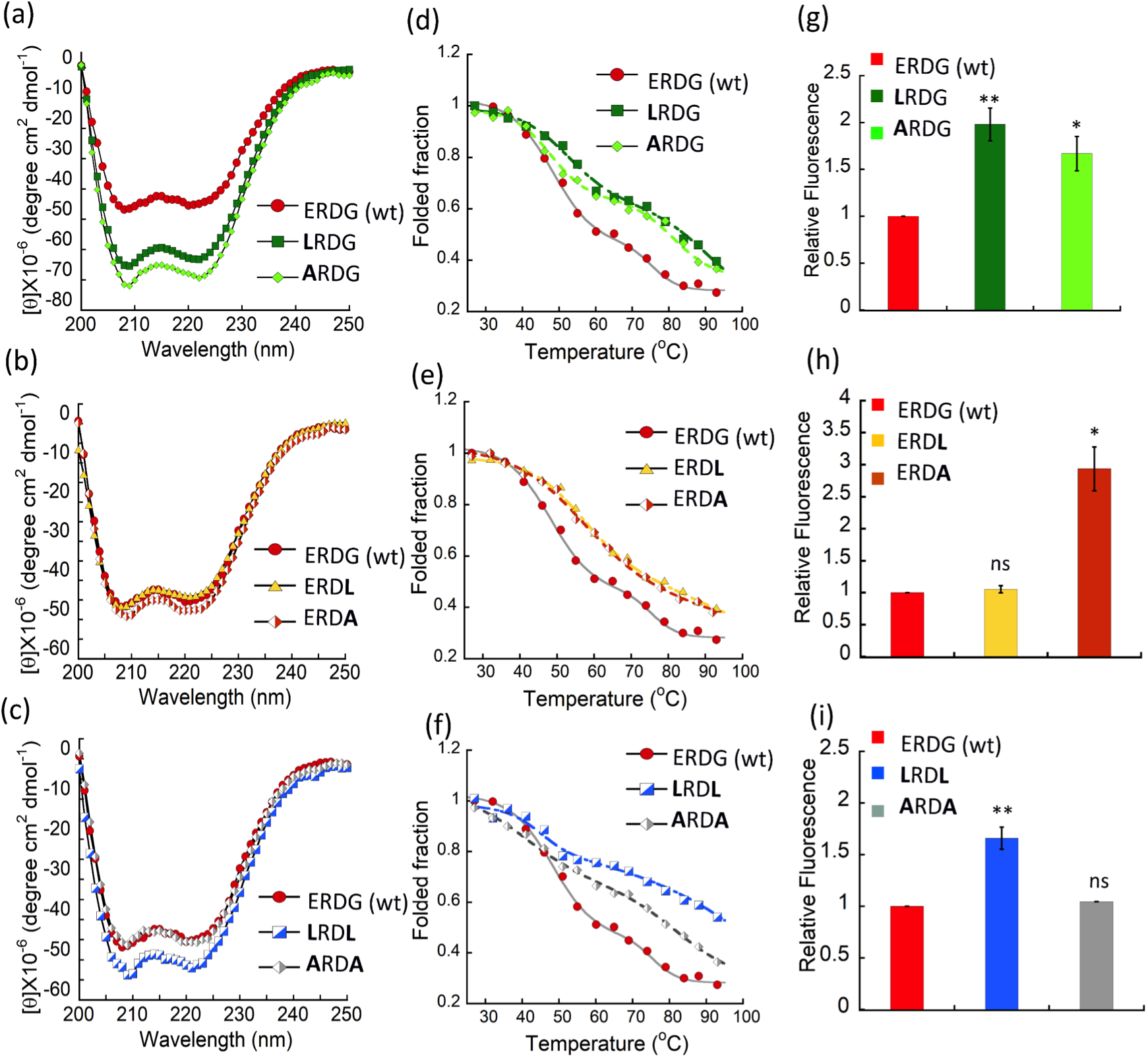
Structural changes in wt-GrpE and stutter mutants. CD spectra of ERDG (wt) (red circle) compared with single stutter mutants **L**RDG and **A**RDG (a), ERD**L** and ERD**A** (b); and double mutants **L**RD**L** and **A**RD**A** (c). Protein concentration was 1 μM in each case. Temperature dependent changes in the folded fraction of ERDG (wt) compared with single stutter mutants **L**RDG and **A**RDG (d), ERD**L** and ERD**A** (e); and double mutants **L**RD**L** and **A**RD**A** (f). Change in intrinsic Tyr-fluorescence for **L**RDG and **A**RDG (g), ERD**L** and ERD**A** (h); and double mutants **L**RD**L** and **A**RD**A** (i). An excitation wavelength of 275 nm was used, and relative fluorescence emission intensity at 303 nm was plotted for 5 μM of ERDG (wt) and all stutter mutants. Here, *p < 0.05, **p < 0.01, ns=non-significant (p > 0.05), as determined from a student t-test for paired data using Kaleidagraph, n=3 for each condition.

To evaluate differences in the stability of these mutants, we performed thermal unfolding experiments for all the mutants. While the **A**RDG mutant did not show any significant difference in T_m1_, **L**RDG showed an increase of ~5°C indicating that introduction of leucine stabilizes the coiled coil domain. More interestingly, T_m2_ increased significantly for both **L**RDG and **A**RDG mutants (~15°C and ~6°C respectively) (Fig. 3d). As the second melting transition has been attributed to unfolding of the four helix bundle^15^, the thermal denaturation data suggests that the effect of introducing a single hydrophobic residue has propagated to the core, and significantly increased the stability of the four helical bundle region. Surprisingly, for mutants at position 61, i.e. ERD**L** and ERD**A**, we found only one transition with an intermediate T_m_ value approx. 57°C (Fig.3e), suggesting that the glycine residue at this position might be important in cross-talk between these two domains of the protein. In contrast, the double stutter mutants **L**RD**L** and **A**RD**A** (Fig. 3f) showed a decrease in T_m1_ by ~4°C and 8°C respectively, but a notable increase in T_m2_ for both the double mutants. While **A**RD**A** showed a similar increase in T_m2_ as the **A**RDG mutant, the **L**RD**L** mutant was so stable that we were unable to determine the second transition for **L**RD**L** in the given experiment setup (Fig. 3f, Table 1). These results indicate that introduction of hydrophobic mutations in the stutter leads to more stable interhelical interactions, and increase in overall protein stability. In particular, the double stutter mutant **L**RD**L** resulted in a greatly increased thermal stability of the four helical bundle region, ~45Å away from the mutation sites.

### GrpE undergoes a conformational change due to a more hydrophobic stutter

To check whether the stutter mutations induced any significant conformational changes in GrpE, we used a tyrosine fluorescence assay. Each GrpE monomer contains one Tyr residue between the core of the β-sheet and the four-helix bundle. A significant change in protein conformation near the four helix bundle due to mutation in the coiled coil might alter the environment of tyrosine, leading to a change in its fluorescence emission properties. To test this idea, we measured Tyr fluorescence for wt-GrpE and the six stutter mutants.

We observed a significant increase in the fluorescence emission for **L**RDG, **A**RDG and **L**RD**L** mutants (Fig. 3g, i). In comparison, mutation at position 61 (Gly) with Ala seems to rescue any changes due to A at position 58, such that **A**RD**A** has no net change from wt-GrpE. In contrast, we found a significant increase in fluorescence for ERD**A**, and no change in ERD**L** (Fig. 3h). These results support a model where mutation at position 58 (Glu) has propagated to and resulted in a conformational change in the four-helix bundle, whereas the effect of the mutation at position 61 (Gly) is more localized. These results were also supported by the predicted hydropathy index using the ProtScale tool (from ExPaSy-Bioinformatics tool portal)^41^. We observed a significant increase in the hydropathy index for mutants **L**RDG, **A**RDG, **L**RD**L** and **A**RD**A** with increased net hydrophobicity of nearby residues. In comparison to this, a small local change was observed for the ERD**A** and ERD**L** mutants.

However, these results were somewhat different from the data obtained from an *in vitro* ANS-binding assay. ANS is a commonly used dye to probe the surface hydrophobicity of proteins, which usually causes enhanced fluorescence emission of ANS and a blue shift of the peak maxima^42^. We did not observe any significant change in ANS fluorescence for the stutter mutants except **L**RDG, which showed a 66% increase in fluorescence intensity and a prominent blue shift (10 nm) in ANS fluorescence emission. This suggests that although there are local changes in the hydrophobic environment of the protein, the overall hydrophobicity remains unchanged due to mutations. Taken together, the biophysical analysis suggest that Leu or Ala substitution at position 58 in the stutter resulted in more significant alterations in GrpE structure and stability.

### Increased hydrophobicity in the stutter alters DnaK-GrpE interaction

To gain insight into the molecular interactions of GrpE stutter mutants with DnaK, we designed and developed a FRET-based assay to monitor the DnaK-GrpE interaction. The residues suitable for FRET analysis were selected from the structure of GrpE bound with the nucleotide binding domain of DnaK (PDB ID: 1DKG). Ser101 was selected in the four helical bundle region of GrpE for donor labeling, and Glu272 was selected in DnaK-NBD for acceptor labeling. The distance calculated from the crystal structure between the two residues was 11.4 Å, which makes it a potentially good FRET pair (Fig. 4a). The native residues in both proteins were first mutated to Cys, and subsequently labeled with the maleimide derivatives of specified dyes (Donor=DACM, acceptor=BODIPY). We observed no significant change in the functionality of these mutants after cysteine introduction, as confirmed by an ATPase assay. In the FRET experiment, when BODIPY labeled DnaK was added to DACM labeled GrpE, we observed a ~75% decrease in donor fluorescence with a corresponding increase in acceptor fluorescence (Fig. 4b). The FRET signal could be chased with unlabeled GrpE, indicating that the FRET signal was indeed reporting on the DnaK-GrpE interaction. The FRET signal could also be chased with ATP and ADP, which are biological effectors for the dissociation of the DnaK-GrpE complex in the DnaK chaperone cycle.

**Fig. 4.**
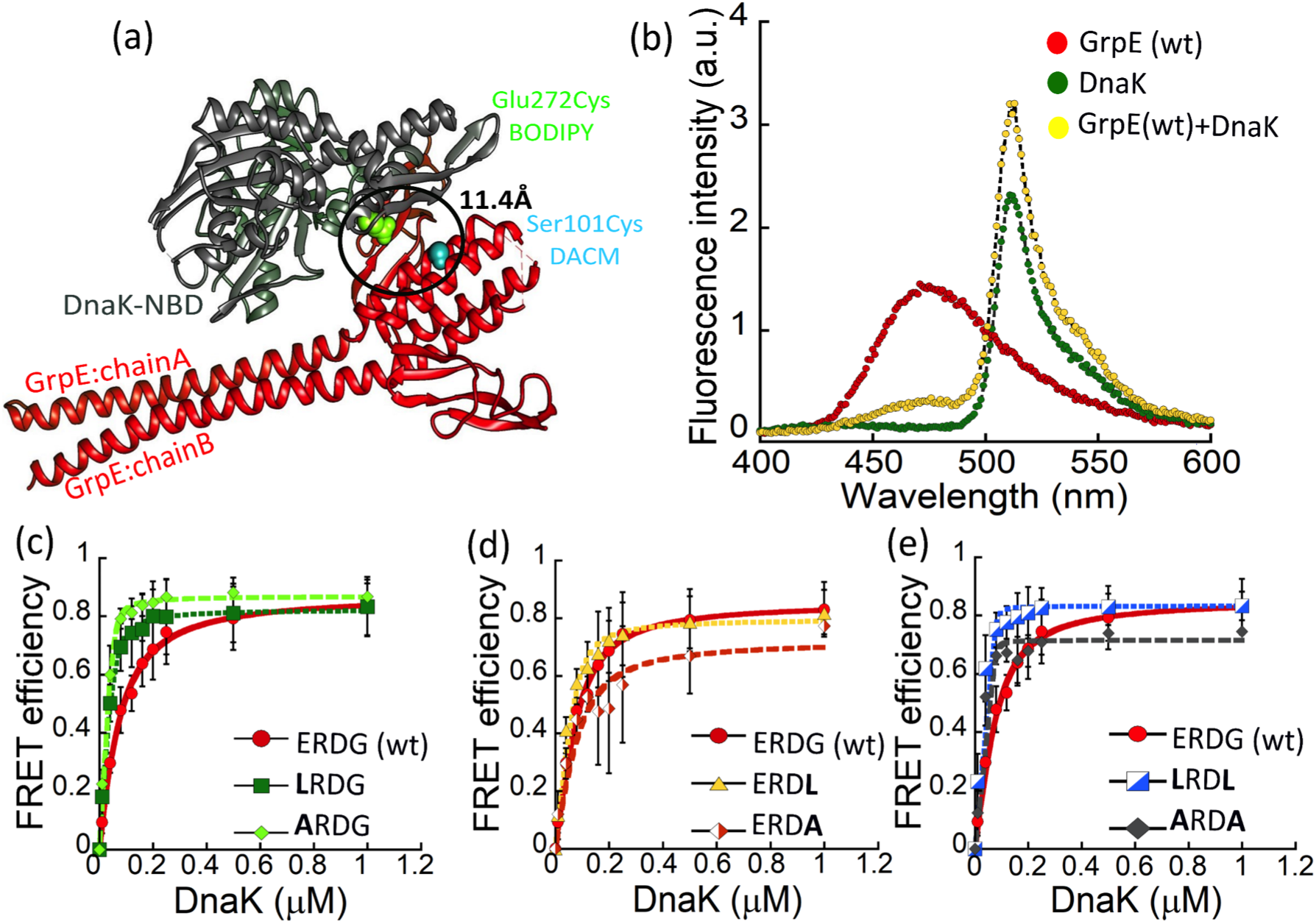
*In vitro* FRET assay between DnaK and GrpE (wt and mutants). (a) The positions of the amino acid residues selected for the DnaK-GrpE FRET assay are depicted in the co-crystal structure (PDB ID: 1DKG). GrpE was labeled at position 101 with DACM as FRET donor, and DnaK was labeled at position 272 with BODIPY as FRET acceptor. Using the crystal structure, the distance between the two residues was estimated to be 11.4 Å. (b) Plot showing fluorescence of donor labeled 0.1 μM GrpE(wt), acceptor labeled 0.2 μM DnaK, and a mixture of the two. Excitation wavelength = 380 nm. Plots showing equilibrium titrations between GrpE or its mutants (0.1 μM) and DnaK, for mutants **L**RDG and **A**RDG (c), ERD**L** and ERD**A** (d), **L**RD**L** and **A**RD**A** (e).

To check whether mutations in the stutter impacted the binding affinity of GrpE with DnaK, we titrated acceptor-labeled DnaK with donor-labeled GrpE(wt) and the labeled stutter mutants. The K_d_ (~32 nM) value calculated for the wt-type system was in excellent agreement with previous reports^18,21^. The single stutter mutants **L**RDG and **A**RDG bound to DnaK more tightly with ~20 fold reduction in K_d_ (Fig. 4c), while a larger reduction in the Kd values was observed for the double stutter mutants **L**RD**L** and **A**RD**A** (~ 50 and ~30 fold respectively) (Fig. 4e, Table 2). In contrast to these results, the single stutter mutants at position 61 (Gly) ERD**L** and ERD**A** did not show a significant change in the binding affinity (Fig. 4d, Table 2). These results are consistent with the biophysical analyses described in the earlier sections. Collectively, our data strongly imply that mutation at position 58 (Glu) of the stutter leads to a significant change in the DnaK-GrpE binding interaction, and enhances the binding affinity between GrpE and DnaK.

**Table 2.** K_d_ values for GrpE mutants with DnaK, compared to wt-GrpE.

|  | ERDG (wt) | LRDG | ARDG | ERDL | ERDA | LRDL | ARDA |
| --- | --- | --- | --- | --- | --- | --- | --- |
| $K_d$ (nM) | $32.8 \pm 4$ | $1.8 \pm 0.2$ | $1.6 \pm 1.5$ | $10.8 \pm 1.1$ | $23.7 \pm 2.3$ | $0.6 \pm 0.4$ | $0.9 \pm 0.8$ |
| fold change | - | 18 | 20 | 3 | 1.4 | 54 | 36 |

### Mutation in the stutter modulates the ATPase and substrate refolding activity of DnaK

The co-chaperonic function of GrpE activates nucleotide exchange in DnaK. To check whether and how GrpE stutter mutants modulate the ATPase activity of DnaK, we performed a malachite-green based colorimetric assay (Fig. 5a). This assay measures the amount of inorganic phosphate released during ATP hydrolysis by DnaK, and is often used as a readout for ATPase activity^43^. Most of the mutants had, at best, a modest effect on the DnaK ATPase activity. We observed a ~ 40% inhibition of DnaK ATPase activity for the double stutter mutant **A**RD**A** (Fig. 5b), while for the ERD**A** and **L**RD**L** mutants we observed a 30% increase in ATPase activity. No significant change was observed for the other three stutter mutants (Fig. 5b). Although the biophysical assays showed significant conformational changes due to increase in hydrophobicity at position 58, we find that the ATPase activity of DnaK is minimally affected. This result suggests that the coiled coil region is functionally decoupled from the residues of GrpE involved in opening of the DnaK nucleotide binding pocket^15^.

**Fig. 5.**
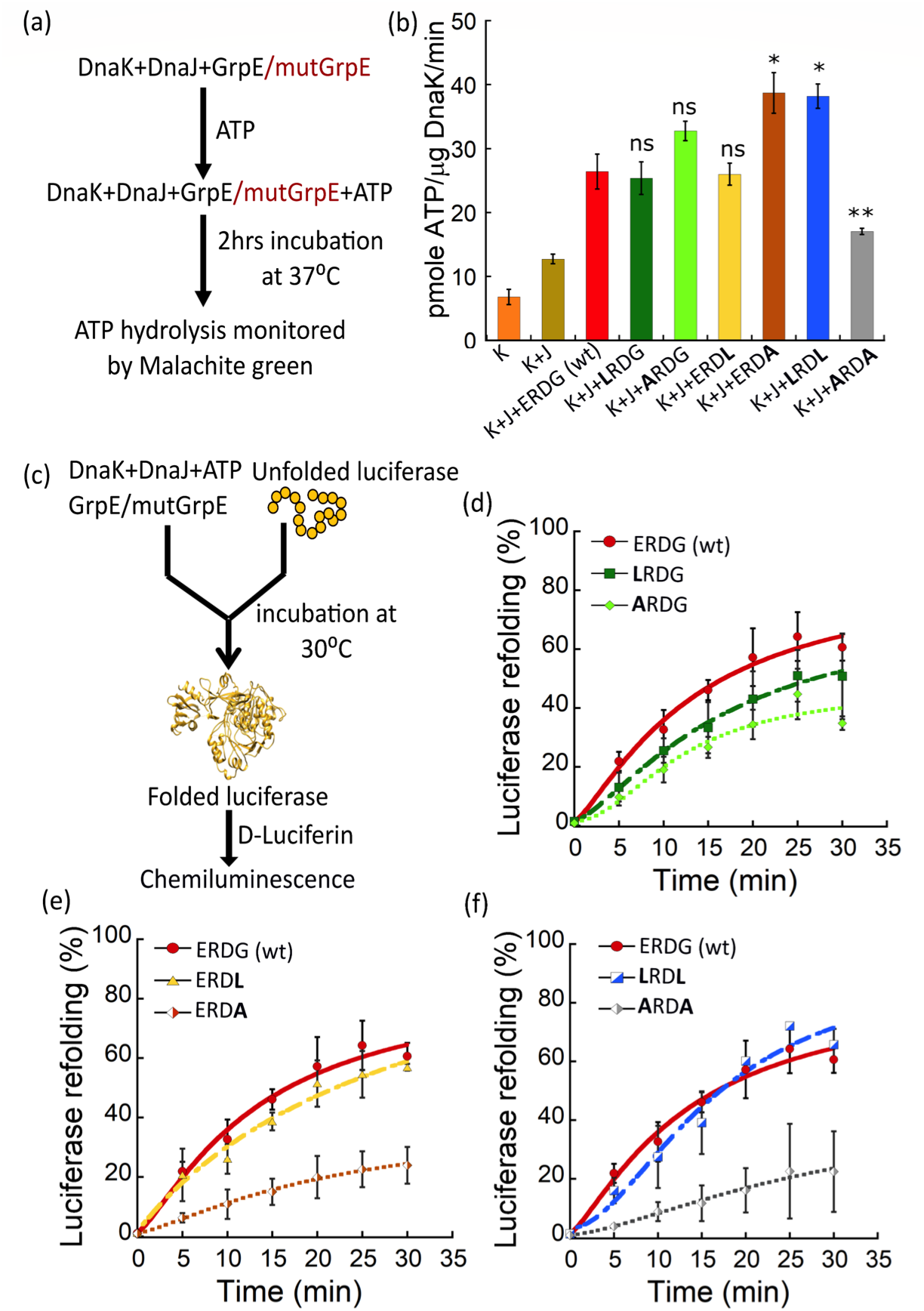
*In vitro* functional assays for GrpE stutter mutants. (a) Schematic illustration of the *in vitro* ATPase activity assay. Plots shows ATPase activity for 0.5 μM ERDG (wt) (red bar) and 0.5 μM single stutter mutants: **L**RDG (green), **A**RDG (light green), ERD**L** (yellow) and ERD**A** (brown); and double stutter mutants: **L**RD**L** (blue) and **A**RD**A** (grey). [ATP] = 1 mM; [DnaK] = 0.7 μM (K) and [DnaJ] = 0.5 μM (J). Here, *p < 0.05, **p < 0.01, ns=non-significant (p > 0.05), as determined by a student t-test for paired data using Kaleidagraph, n=3 for each case. (c) Schematic illustration of the *in vitro* luciferase refolding assay. (d-f) Plots showing the relative folding of chemically denatured 0.08 μM luciferase in the presence of 2 mM ATP, 0.8 μM DnaK, 0.16 μM DnaJ, and 0.4 μM ERDG (wt) (red) or 0.4 μM stutter mutants **L**RDG and **A**RDG (d); ERD**L** and ERD**A** (e); and **L**RD**L** and **A**RD**A** (f). Luciferase activity was normalized to the activity of equal concentration of native luciferase.

The other important co-chaperonic activity of GrpE is to release folded substrate proteins from DnaK to ensure efficient folding turnover. To test the protein folding efficiency with GrpE mutants, we performed a luciferase refolding assay. In this assay, the percentage of refolded luciferase is quantified by measuring the luminescence emitted by fully folded luciferase in the presence of its substrate D-luciferin (Fig. 5c). We observed a significant reduction in luciferase refolding activity for both mutants containing alanine at position 61: ERD**A** and **A**RD**A**, showing 40-60% reduction in refolding activity (Fig. 5e, f). We also observed somewhat reduced refolding activity for **L**RDG and **A**RDG (~20%) (Fig. 5d), whereas no significant change was observed for ERD**L** and **L**RD**L** mutants (Fig. 5e, f). The reduction in refolding activity for both the Gly61Ala mutants indicates that these mutations likely affected the flexibility of the N-terminal domain on the other end of the coiled coil, away from the ATPase binding site. Our results are in agreement with previous reports suggesting the functional importance of the N-terminal portion of GrpE in substrate refolding^21,44,45^.

### Altered structural stability of GrpE coiled-coil affects bacterial cell growth

GrpE is essential during heat stress as it acts as a thermosensor, and regulates the growth of bacterial cells. To test the effect of the GrpE stutter mutants during bacterial growth, we performed *in vivo* complementation studies on a *grpE* deleted strain. For this experiment, we used a *grpE*-deleted *E. coli* DA259 strain^46^ and complemented it with a plasmid containing either wt-GrpE or stutter mutants. The cells were grown under two different temperature conditions 30°C and 42°C (Fig. 6a,b). All cells grew normally at 30°C, as GrpE deletion is not lethal at that temperature^10^. At heat shock temperature (42°C), DA259 cells complemented with double mutants **L**RD**L** or **A**RD**A** showed significant defect in growth after25 hrs (Fig. 6b). The extent of defect was comparatively smaller for the single stutter mutants. This result indicates that during heat stress the enhanced binding affinity between DnaK and GrpE is detrimental to bacterial growth, likely because it disrupts interconversion of DnaK between its foldase to holdase states, and prevents protein folding. Taken together, these results highlight that both stability and flexibility in the coiled-coil domain of GrpE are crucial in controlling its thermosensing activity.

**Fig. 6.**
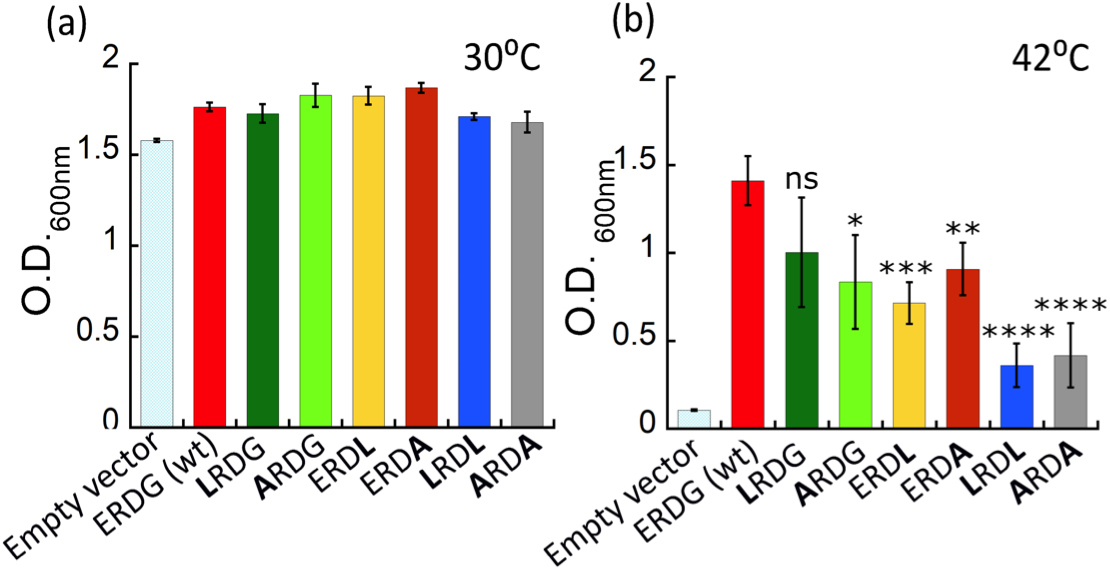
*In vivo* growth assays for stutter mutants. Bacterial growth assays for *E. coli* DA259 cells complemented with ERDG (wt) (red bar); single stutter mutants: **L**RDG (green), **A**RDG (light green), ERD**L** (yellow) and ERD**A** (brown); and double stutter mutants: **L**RD**L** (blue) and **A**RD**A** (grey) at (a) 30°C (after 20 hrs) and (b) 42°C (after 25 hrs). *p < 0.05, **p < 0.01, ***p < 0.001, ****p < 0.0001, ns=non-significant (p > 0.05), as determined by a student t-test for paired data using Kaleidagraph, n=6 for each case.

## Discussion

There are several examples of proteins having non-heptad periodicities in the coiled-coil domain, but only a few studies have reported their functional significance^40,47,48^. In this work, we have explored the role of an extended helical region in *E. coli* co-chaperone GrpE, which forms a long non-ideal coiled-coil in the GrpE dimer. The coiled-coil plays a crucial role in the thermosensing activity of GrpE and in regulating its structure and function^16^. *E. coli* GrpE displays two well-separated thermal transitions with midpoints at ~50°C and ~75°C which have been attributed to the unfolding of the coiled-coil domain, and the four helical bundle domain respectively. During heat stress, the long coiled-coil of GrpE undergoes significant structural changes, and plays an essential role in sensing the change in ambient temperature to activate the heat shock response.

In this work, we propose that deviation from an ideal coiled-coil domain has evolved as an important structural feature of GrpE to tailor survival of the organism at the requisite temperature. This deviation is achieved through introduction of stutters in the coiled-coil domain, which increases or decreases the coiled coil probability, and controls the structure-function relationship. However, the molecular mechanism by which the stutter in the GrpE coiled-coil domain influenced its thermosensing activity and whether it impacted the interaction of GrpE with DnaK was not known. In this work, we have studied how changes in the stutter regulate the structural stability of GrpE, and its impact on the DnaK chaperone cycle.

We performed a mutational analysis at the first stutter of the *E. coli* GrpE coiled-coil domain comprising “ERDG” residues, having a destabilizing residue (Glu) in the hydrophobic core, and a helix breaker (Gly) at the end of the stutter. We mutated Glu58 or Gly61 in the stutter to Leu or Ala residues, and monitored their impact on stability and function of GrpE. Interestingly, substituting Leu or Ala at position 61 of the coiled-coil domain to generate mutants ERD**L** and ERD**A** resulted in an increase in overall protein thermal stability, and merging of the two thermal melting transitions into one. The thermal unfolding patterns for these mutants were similar to yeast GrpE (Mge1p) and human mitochondrial GrpE (GrpE#1) both of which have only one transition temperature^49,50^. A multiple sequence alignment (MSA) study comparing GrpE sequences from several organisms showed that glycine is present at that location only in *E. coli* GrpE. We speculate that given the very low helix propensity of glycine, its presence in *E. coli* GrpE allows structural flexibility and tuning of co-chaperone function. Thus, it is likely that Gly at position 61 maintains the conformational flexibility of the protein, and mutation at this position results in heat sensitivity during bacterial growth.

Similarly, introducing a hydrophobic residue in the stutter either at position 58, or both at positions 58 and 61 strengthened the hydrophobic core due to increased interhelical interaction interface (Fig. 2 c-d, g-h). We note that all single mutants having a mutation at position 58 showed a significant increase in T_m2_, although the double mutants showed a modest decrease in T_m1_. This observation indicates that mutation to leucine at position 58 of the coiled-coil domain results in the structural rearrangements that are propagated towards the four helical bundle region, and increase the stability of that region (Table. 1).

Previously it has been reported that increased stability of a coiled-coil limits its conformational space, which alters its specific protein binding interfaces^47^. GrpE has a long extended structure which undergoes several conformational changes during thermal stress, and during its interaction with DnaK^16^. Similarly, DnaK shows numerous conformational changes during its chaperone activity, exposing protein binding interfaces for interaction with its partner proteins DnaJ and GrpE. To test whether the altered stability of the coiled coil affects the GrpE-DnaK interaction, we developed a FRET assay to study this protein-protein interaction. The FRET assay was validated by determining the binding affinity between wild-type DnaK-GrpE with a K_d_ of ~32 nM, in agreement with previous reports^18^.

We found that the increased GrpE stability in the four helical bundle region due to mutations in the stutter translated into an increased affinity for DnaK. Removal of Glu from position 58 resulted in mutants that bound more tightly with DnaK, consistent with our earlier result showing increased stability of the four helical bundle in these mutants. It is plausible that since position 58 in the stutter is facing the core of the coiled-coil domain, substituting hydrophobic residues may lead to either structural rearrangements or interhelical collapse. The thermal melting data showed increased stability in the four helical bundle region, suggesting that structural rearrangements in these mutants resulted in an increased affinity for DnaK. The single stutter mutants **L**RDG and **A**RDG showed ~20 fold increase in affinity, and double stutter mutants **L**RD**L** and **A**RD**A** showed ~50 and 30 fold increase in affinity respectively. It has been proposed that during DnaK-GrpE interaction, the head region of GrpE interacts first with DnaK-NBD at the designated hydrophobic patches on the interface. This is followed by the interaction of the long helical region with the DnaK linker domain, and finally the first 33-residues of GrpE act as a pseudo-substrate, interacting with the substrate binding pocket of DnaK-SBD^44,45,51^. It is plausible that the increase in DnaK-GrpE binding affinity is due to selection of a preferred GrpE conformation, or because the conformational changes at the binding interface result in more favorable binding interactions between the two proteins. As the co-crystal structure (PDB ID: 1DKG) neither contains the DnaK substrate-binding domain nor the DnaK interdomain linker which are involved in interacting with the coiled-coil domain of GrpE^18^, our basic structural knowledge of the interaction is limited^52^. This study results in an improved understanding of the intermolecular communication between the two proteins.

The DnaK-GrpE interaction is highly dynamic, and a small change in affinity may affect the dynamic behaviour of these interacting partners. The structural changes due to GrpE stutter mutations were not manifested in the ATPase activity of DnaK, either because the changes were too small or did not impact the relevant portions of GrpE^53^. This effect of the altered DnaK-GrpE interaction was more evident in the luciferase refolding assay, where mutation with Ala at position 61, showed a significant decrease in luciferase refolding activity. This result indicates more structural changes towards the N-terminal region of GrpE on the other end of the coiled-coil domain, affecting the interaction interface of GrpE with DnaK-SBD.

The defect in co-chaperonic activity was further confirmed by a bacterial cell growth assay, where most of the mutGrpE complemented strains showed intolerance to heat shock temperature (42°C)^10,45,52^. These results further suggest that mutant proteins showing tighter binding affinity between DnaK and GrpE could hold the DnaK-GrpE complex in one particular state (holdase state), disrupting the chaperonic function during heat stress^54,55^. It has also been reported that GrpE has a role in stimulating the ATPase activity of Hsc62 (third Hsp70 homolog) in *E. coli* in the presence of Hsc56^56^. Thus, it is possible that mutant GrpE proteins differentially affect both DnaK and Hsc62 chaperone systems inside the cell.

In summary, we propose a model in which a stutter in the coiled-coil domain of GrpE is responsible for providing flexibility to the protein (Fig. 7). This flexibility is crucial for interaction with DnaK, and to regulate its conformational equilibrium^16,45^, and eventually control chaperone function. Mutation in the stutter region with hydrophobic residues modulates the structural stability of the protein, and affects the conformational dynamics of GrpE leading to a defect in the DnaK chaperone cycle. This study gives critical insights into the role of a stutter in the coiled-coil domain of GrpE, and its involvement in conformational rearrangements during heat stress. Further, it will be interesting to study the role of the second stutter in the structural stability of GrpE, and its functional role in the DnaK chaperone pathway. Overall, this study enhances the understanding of sequence specificity in one stutter of the coiled-coil domain of GrpE, and sets the stage to probe the functional significance of non-ideality in other coiled coils. Our work will be also useful to explore GrpE as a potential antibacterial target in pathogenic bacteria like *M. tuberculosis* (Mtb), where GrpE acts as an immune activator and enhances host-pathogen interaction^57^.

**Fig. 7.**
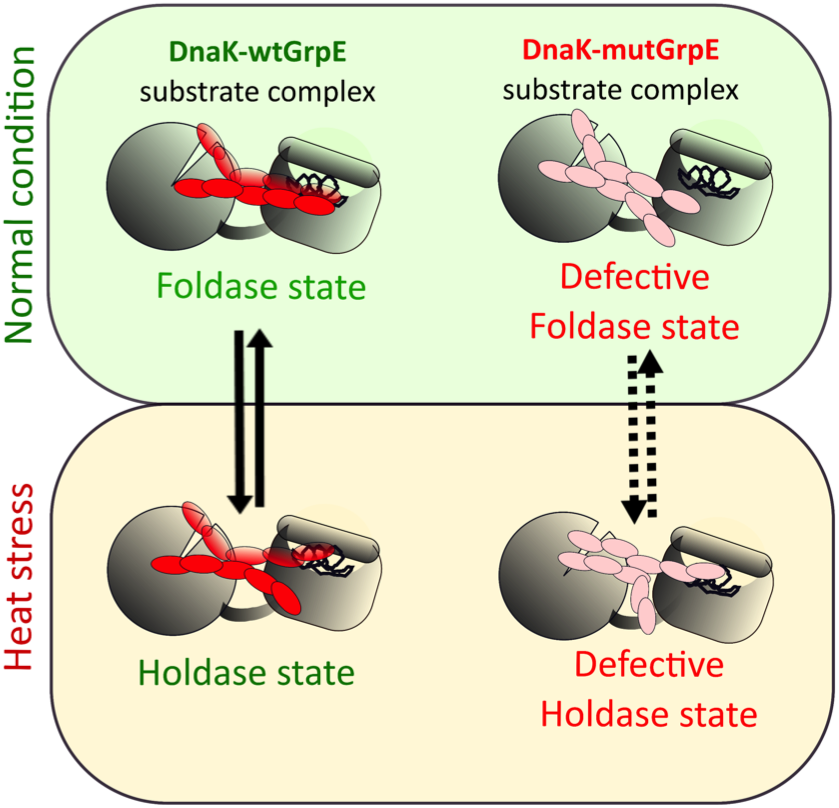
Model for disruption of DnaK chaperone activity by GrpE stutter mutants. wtGrpE (left) regulates the interconversion between the foldase and holdase functions of DnaK to maintain proteostasis. GrpE stutter mutants bind tightly to DnaK, causing a defect in the holdase to foldase interconversion resulting in heat sensitivity of *E. coli*.

## Materials and Methods

### Protein purification

The plasmids for expression of *E. coli* DnaK and GrpE were a kind gift from Bernd Bukau. wtDnaK and wtGrpE, with N-terminal 6His-SUMO, were overexpressed from a pET vector in *E. coli* BL-21(DE3) cells. Cells were grown in LB media supplemented with kanamycin (50 μg/ml) at 30°C till OD reached to 0.7-0.8 (600 nm). Protein expression was induced with 1 mM IPTG and cells were grown overnight at 25°C and purified as described earlier^58^. GrpE stutter mutants **L**RDG, **A**RDG, ERD**L**, ERD**A, L**RD**L** and **A**RD**A** were generated through site-directed mutagenesis at positions E58 and G61 of *E. coli* GrpE and confirmed by DNA sequencing. For purification of all the proteins, cells were harvested by centrifugation and resuspended in lysis and wash buffer, LWB containing 40 mM HEPES (pH 7.5), 150 mM KCl, 5 mM MgCl_2_ 10 mM β-mercaptoethanol, 1 mM PMSF, 0.02% triton-X and 5% glycerol. Cells were lysed by sonication, loaded onto a Ni-NTA column and protein was eluted with 250 mM imidazole. The SUMO tag was cleaved by ULP1 protease by incubating overnight in dialysis buffer and passed through Ni-NTA to get rid of 6His-SUMO. The proteins were further purified using a MonoQ 10/100 column (GE Healthcare) and eluted with a KCl salt gradient from 100 mM to 600 mM. The protein containing fractions were concentrated in an Amicon filter (Millipore) with cut off of 10 kDa (GrpE) and 30 kDa (DnaK). Proteins were stored in 50% glycerol at -30°C. The concentration of protein was measured by Bradford assay with BSA as a standard protein.

DnaJ was overexpressed and purified in BB2583 strains (kind gift from Bernd Bukau) as described elsewhere^59^. Cells were grown in 2xYT medium supplemented with 0.4% dextrose, ampicillin (100 μg/ml) and kanamycin (50 μg/ml) at 30°C till OD reached to 0.3-0.4 (600 nm). Protein expression was induced with 0.5mM IPTG and the cells were grown at 30°C for an additional 5hrs. Cells were harvested by centrifugation and resuspended in lysis buffer (50 mM Tris-HCl (pH 8), 10 mM DTT, 0.8% (w/v) Brij 58, 1 mM PMSF and 0.8 mg/ml lysozyme). Cells were lysed by sonication, and the protein was precipitated with ammonium sulphate. The precipitated protein was dissolved in wash buffer (50 mM sodium phosphate pH 7.0, 5 mM DTT, 1 mM EDTA, 0.1% (w/v) Brij 58) and dialyzed for 4 hrs. The dialyzed protein was purified using a SP-Sepharose column (GE Healthcare), and eluted with a KCl salt gradient (100 mM to 1M). The purified fractions were dialyzed overnight against dialysis buffer (50 mM Tris-HCl (pH 7.5), 5 mM DTT, 0.1% (w/v) Brij 58, and 50 mM KCl). The protein was concentrated in a 10 kDa Amicon filter (Millipore), and purified on a CHT Ceramic Hydroxyapatite Type II (Biorad) column, with a linear gradient of 0 to 50% elution buffer (50 mM Tris-HCl (pH 7.5), 5 mM DTT, 0.1% (w/v) Brij 58, 50 mM KCl and 600 mM KH2PO4). The purified fractions were pooled, dialyzed (50 mM Tris-HCl (pH 7.7 and 100 mM KCl), concentrated, and stored in 50% glycerol at -30°C. The concentrations of purified protein were determined by a Bradford assay

Firefly luciferase was expressed in an *E. coli* strain containing two plasmids pDS56-Luci (selection ampicillin) and pLacIq (selection spectinomycin) obtained from Chandan Sahi (originally from Bukau Lab)^59^. Cells were grown in LB medium supplemented with ampicillin (100 μg/ml) and spectinomycin (50 μg/ml)) at 30°C until OD_600_ = 0.6 was reached, at which point the temperature was lowered to 20°C. After shaking at 20°C for 45 min, cells were induced with 0.1mM IPTG overnight. The cell pellet was resuspended in pre-cooled lysis buffer (50 mM sodium phosphate pH 8.0, 300 mM NaCl, 10 mM β-mercaptoethanol, 1 mM PMSF). Cells were lysed by sonication, and the lysate was loaded onto a Ni-NTA column. The protein was eluted with buffer containing 250 mM imidazole. Purified luciferase was dialyzed overnight in dialysis buffer (50 mM sodium phosphate pH 8.0, 300 mM NaCl and 10 mM β-mercaptoethanol, 10% glycerol). The dialysed protein was concentrated before flash freezing in liquid N2 and stored at -80°C. The concentrations of purified protein were determined by a Bradford assay.

### Protein labeling

For protein labeling, Cys-mutants were generated by site directed mutagenesis at DnaK E272C and GrpE S101C and verified by DNA sequencing. To ensure specific labeling at cysteine 272 in DnaK, the native cysteine in DnaK at residue 15 was mutated to alanine. The Cys-variants for DnaK (DnaK-E272C) and GrpE (ERDG-S101C, **L**RDG-S101C, **A**RDG-S101C, ERD**L**-S101C, ERD**A**-S101C**, L**RD**L**-S101C and **A**RD**A**-S101C) were purified as described for the wild type proteins. All GrpE variants were labeled with DACM (N-(7-dimethylamino-4-methylcoumarin-3-yl)maleimide) (AnaSpec) and DnaK was labeled with BODIPY-fluorescein-N-(2-aminoethyl)-maleimide (Invitrogen) using maleimide-thiol chemistry as described^60^. For DnaK, protein was dialysed against labeling buffer (50 mM HEPES (pH 7.0), 300 mM NaCl, 2 mM EDTA and 5% glycerol) for 2 hrs, followed by incubation with 2 mM TCEP. Then 5-fold molar excess of dye was added, and incubated for 4 hrs at 4°C. The unreacted dye was removed by gel filtration chromatography using Sephadex G-25 resin (GE Healthcare). For GrpE, labeling buffer of pH 8.0 was used and DACM dye was added at 15-fold molar excess. The amount of labeled protein was determined using the extinction coefficient of each dye (ε_383_ = 27,000 M^-1^ cm^-1^ for DACM and ε_504_ = 80,000 M^-1^ cm^-1^ for BODIPY). The concentration of the total protein was determined by Bradford assay. The efficiency of protein labeling was typically ≥ 95% for DnaK, and ≥ 50% for wtGrpE and mutGrpE proteins.

### Steady-state fluorescence and FRET measurements

All steady-state fluorescence measurements were performed at 25°C on Fluorolog-3 (HORIBA Jobin Yvon, Germany). To monitor the binding affinity of wt or mutant GrpE with DnaK, 100 nM GrpE in FRET buffer (100 mM HEPES pH 7.4, 20 mM KCl, and 6 mM MgCl_2_, 0.02% Triton X-100) was titrated with DnaK, and the donor fluorescence was measured at 470 nm (excitation = 380 nm). FRET efficiency was calculated from fluorescence intensities of donor in presence and absence of acceptor using equation 1, and the data were plotted using Kaleidagraph (Synergy, USA):

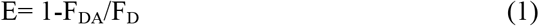

Further, FRET efficiency values were plotted against DnaK concentration and data were fit according to equation 2:

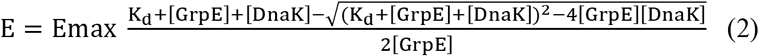

E_max_ is the maximum FRET efficiency at saturating DnaK concentration, and Kd is the equilibrium dissociation constant of the GrpE-DnaK complex.

### Circular dichroism (CD)

CD spectra were recorded using JASCO, J-815 CD spectrophotometer (Jasco, Japan) equipped with a Peltier temperature controller. Single CD spectra were recorded in the far ultraviolet range (200-250 nm, with scan rate: 50 nm/s, band width: 1 nm, data pitch: 1nm. For CD measurements, 1 μM of protein was used in 25 mM sodium phosphate buffer, pH 7.0. Each spectrum is the average of 3 scans corrected by the buffer signal scan and repeated 3 times. The percentage helicity of proteins was calculated using the CD spectra with the tool BeStSel^61^. Thermal melting curves were recorded by monitoring ellipticity at 222 nm at indicated temperature range applying ramp rate of 1°C /min. The data were analysed using Spectra Manager analysis software, version 2 (Jasco, Japan). The data were plotted using Kaleidagraph software (Synergy, USA), and T_m1_ and T_m2_ were estimated from double sigmoidal fits to thermal denaturation plots at 222 nm.

### Colorimetric determination of ATPase activity

The malachite green based assay was performed according to previous reports with slight modifications^43^. Stock solutions of malachite green (Sigma-Aldrich) (0.081% w/v in 6 M H_2_SO_4_), polyvinyl alcohol (Sigma-Aldrich) (2. 3% w/v) and ammonium heptamolybdate tetrahydrate (Merck) (5.7% w/v in 6 M HCl) were mixed with water in ratio of 2:1:1:2 to make malachite reagent mix. To check ATPase activity, 50 μL of reaction mix containing DnaK: DnaJ: GrpE (0.7: 0.5: 0.5 μM) and 1 mM ATP were mixed in ATPase assay buffer (100 mM Tris–HCl pH 7.4, 20 mM KCl, and 6 mM MgCl_2_, 0.02% Triton X-100). After incubation for 2 hrs at 37°C, 160 μL of malachite reagent mix was added in each well, followed by addition of 20 μL of 34% sodium citrate (Sigma-Aldrich) to quench the reaction^43^. Samples were mixed thoroughly and incubated for 15 min at 37°C before measuring absorbance at 620 nm on a BioTek Cytation 1 plate reader. The background signal from ATP alone (without any chaperones) was subtracted from each data point.

### Luciferase refolding assay

Luciferase (10 μM) was chemically denatured by incubating in unfolding buffer (5 M GdnCl, 30 mM Tris-Cl pH 7.5) for 10 mins at 22°C. The denatured luciferase was diluted with refolding buffer (25 mM HEPES-KOH pH 7.6, 100 mM KCl, 10 mM MgCl_2_, 2 mM ATP, 5 mM DTT, 0.8 μM DnaK, 0.16 μM DnaJ and 0.4 μM GrpE or mutants) and incubated for refolding at 30°C (80 nM final concentration of luciferase). For measurement of luciferase activity, at indicated times, refolded sample was diluted 125-fold in assay buffer (100 mM K-phosphate buffer pH 7.6, 25 mM Glycylglycine, 100 mM KCl, 15 mM MgCl_2_, 5 mM ATP)^50^. Luminescence was measured immediately after injecting 125 μL of 80 μM D-luciferin (Sigma-Aldrich) mixed with assay buffer (without ATP) on a BioTek Cytation 1 plate reader. Refolding yields were normalized based on luminescence of equal concentration of native luciferase.

### *In vivo* growth curve study

The GrpE knockout strain *E. coli* DA259 (C600 *ΔgrpE::ΩCmR* thr::Tn10) was obtained from Dr Shinya Sugimoto, Japan (strain originally from Georgopoulos lab at Utah). To control GrpE expression, GrpE gene was cloned into a pBAD Vector. DA259 cells were transformed with ERDG (wt) or mutant GrpE in pBAD vector. The primary culture was grown till OD_600_ = 0.2, and growth curves were monitored in 96 well plates (5 × 10^2^ CFU/200μl culture). The expression of protein was induced with 3mM arabinose. The growth data were recorded at 30°C and 42°C, with a BioTek Cytation 1 plate reader for 20-25 hrs.

## Acknowledgment

We thank IISER Bhopal for financial support. I.S is a recipient of Ramanujan Fellowship from the Science and Engineering Research Board. We thank Prof. Bernd Bukau for DnaK, GrpE and luciferase expression plasmids, and DnaJ expression strain. We thank Prof. Costa Georgopoulos and Prof. Shinya Sugimoto for providing *E. coli* GrpE knockout strain DA259, and Prof. Chandan Sahi and Yogesh Tak for providing the luciferase expression construct, and purified luciferase.

